# Detecting spatial dynamics of range expansions with geo-referenced genomewide SNP data and the geographic spectrum of shared alleles

**DOI:** 10.1101/457556

**Authors:** Diego F. Alvarado-Serrano, Michael J. Hickerson

## Abstract

Uncovering the spatial dynamics of range expansions is a major goal in studies of historical demographic inference, with applications ranging from understanding the evolutionary origins of domesticated crops, epidemiology, invasive species, and understanding specieslevel responses to climate change. Following the surge in advances that make explicit use of the spatial distribution of genetic data from georeferenced SNP variants, we present a novel summary statistic vector, the geographic spectrum of shared alleles (GSSA). Using simulations of twodimensional serial expansion, we find that the information from the GSSA, summarized with Harpending’s Raggedness Index (RI), can accurately detect the spatial origins of a range expansion under serial founder models, even with sparse sampling of only ten individuals. When applying to SNP data from two species of the holarctic butterfly genus *Lycaeides*, the suggested origins of expansion are consistent with hindcasts obtained from ecological niche models (ENMs). These results demonstrate the GSSA to be a useful exploratory tool for generating hypotheses of range expansion with genomewide SNP data. Our simulation experiments suggest high performance even with sampling found in studies of nonmodel organisms (one sampled individual per location, no outgroup information, and only 5,000 SNP loci).

## Introduction

Geographic range shifts are a prevalent feature of virtually all species’ histories. They often follow or accompany the origin of species (Gaston 2003), and frequently happen in response to changes in climate, landscape, or biotic opportunities (Konečný *et al.* 2013 Roberts & Hamann 2015). Despite range expansions being a pervasive feature across many species histories, the net outcomes of the interactions between speciesspecific traits and changes in suitable habitats are poorly understood with regards to changes in the geographic distribution of genetic diversity, adaptability, and community structure (Pharo & Zartman 2007 Thuiller *et al.* 2008 Phillips *et al.* 2010). For example, although serialrange expansions are likely a common feature of species histories in the context of past global changes (Excoffier *et al.* 2009), there is still uncertainty about how they affect differentiation and diversification, or how they influence the risk of extinction under future environmental shifts (Charles & Dukes 2008 Fordham *et al.* 2014). Tools that uncover the spatiotemporal dynamics of species’ ranges are therefore critical for understanding how geographic range dynamics affect the spatial distribution of genetic diversity through drift and selection (Slatkin 1987 Kirkpatrick & Barton 1997 Peischl *et al.* 2013). They are also central to forecast species’ responses to ongoing and future environmental changes (Brown *et al.* 2016).

Among recent attempts to couple genetic and spatial information to infer historical range dynamics, Ramachadran *et al.* (2005) proposed to identify the geographic origin of an expansion using heterozygosity clines that result from the reduction of heterozygosity associated with the sequential founder events that are characteristic of range expansions (Ramachandran *et al.* 2005 Li *et al.* 2008 DeGiorgio *et al.* 2009 Henn *et al.* 2012). Based on the premise that heterozygosity should continuously decrease with distance from the expansion source, this method fits a linear regression between expected heterozygosity at multiple sampled locations and their geographic distance from all possible candidate sources. While this is simple to estimate, this approach has been found to perform well under a relatively narrow range of circumstances, with its accuracy decreasing as the speed of the expansion decreases, or as the time since the end of the expansion increases (Peter & Slatkin 2015). Previously, researchers have used principal component analysis (PCA) of georeferenced samples to identify the direction of expansion under the assumption that the first principal component—which accounts for the greatest amount of variation—aligns with the expansion sourcefront axis (Menozzi *et al.* 1978). However, this assumption has been challenged by recent evidence indicating that the geographic alignment of the first component may be driven by the geographic distribution of samples, allele surfing, or mathematical artifacts (Novembre & Stephens 2008 François *et al.* 2010 Pemberton *et al.* 2013).

More recently, Peter and Slatkin (2013) proposed a new statistic, the directionality index (ψ), to detect the origins of range expansions using joint georeferenced and genomewide information. This statistic takes advantage of allele frequency changes driven by the sequential founder events that often characterize expansions. Specifically, ψ detects asymmetries in 2D site frequency spectra of shared neutral nonancestral alleles between pairs of populations, under the theoretical expectation that populations further away from the expansion have experienced more genetic drift than populations near the expansion source (Slatkin & Excoffier 2012 Peter & Slatkin 2013). This metric has been also incorporated into an ABC framework, whereby He et al. (2017) used it to estimate the source of range expansion in the Collared pika in Alaska.

While analyses using the directionality index (Peter & Slatkin 2015) have been highly promising, the relatively high number of sampled individuals, and the requirement of polarized SNPs, make its implementation difficult in non-model organisms, which are typically characterized by limited numbers of georeferenced genotypes, or for which a reliable outgroup for polarizing SNPs has not been identified. Here we develop a summary statistic vector, which we call the geographic spectrum of shared alleles (GSSA, hereafter), to harvest information from the spatial distribution of shared allelic variants to help identify the geographical origin of range expansions. Our approach is targeted for nonmodel organisms as it does not require information about ancestral states, requires a single individual per sampled location, and can accommodate a reasonable number of SNPs.

We formally describe the GSSA summary statistic vector and use two-dimensional simulations of different expansion histories to statistically explore its behavior. We also demonstrate its applicability with genomewide SNP data sampled from a set of taxa of the holartic butterfly genus *Lycaeides* distributed in Western North America (Gompert *et al.* 2010). Our analyses demonstrate that the GSSA can be a powerful exploratory tool to identify the relative position of samples along an expansion axis. In *Lycaeides*, the GSSA places the origin of expansions in geographical areas that are consistent with species distribution models for hypothesized ranges shifts during the last glacial maximum.

## Materials and Methods

### The geographic spectrum of shared alleles (GSSA)

The Geographic Spectrum of Shared Alleles (GSSA) is a property of—and calculated for—each sampled locality. It captures the geographic distribution of shared coancestry with other localities by summarizing the geographic distances between copies of the minor allele observed at each SNP site (that is, the allele with the smallest frequency in the global sample for each site Nielsen et al. 2012). As such, the GSSA of a given location is a histogram that depicts the cumulative frequency distribution of the geographic distances separating each minor allele shared between the focal location and all other sampled locations across all SNPs (Fig. 1). On an intuitive level, this new summary statistic vector can be understood as a spatial structure function that captures the relative change in allelic similarity from a focal location as a function of geographic distance. Importantly, the GSSA is a location-specific statistic which differs from other structure functions such as spatial correlograms that are globally calculated. These latter statistics quantify the spatial dependence of an observed variable across the entire set of observations using a covariance matrix partitioned into distance classes (Smouse & Peakall 1999 Legendre & Legendre 2012). Instead, each location-specific GSSA is directly based on distances between minor alleles shared with a focal locality.

**Fig. 1.**
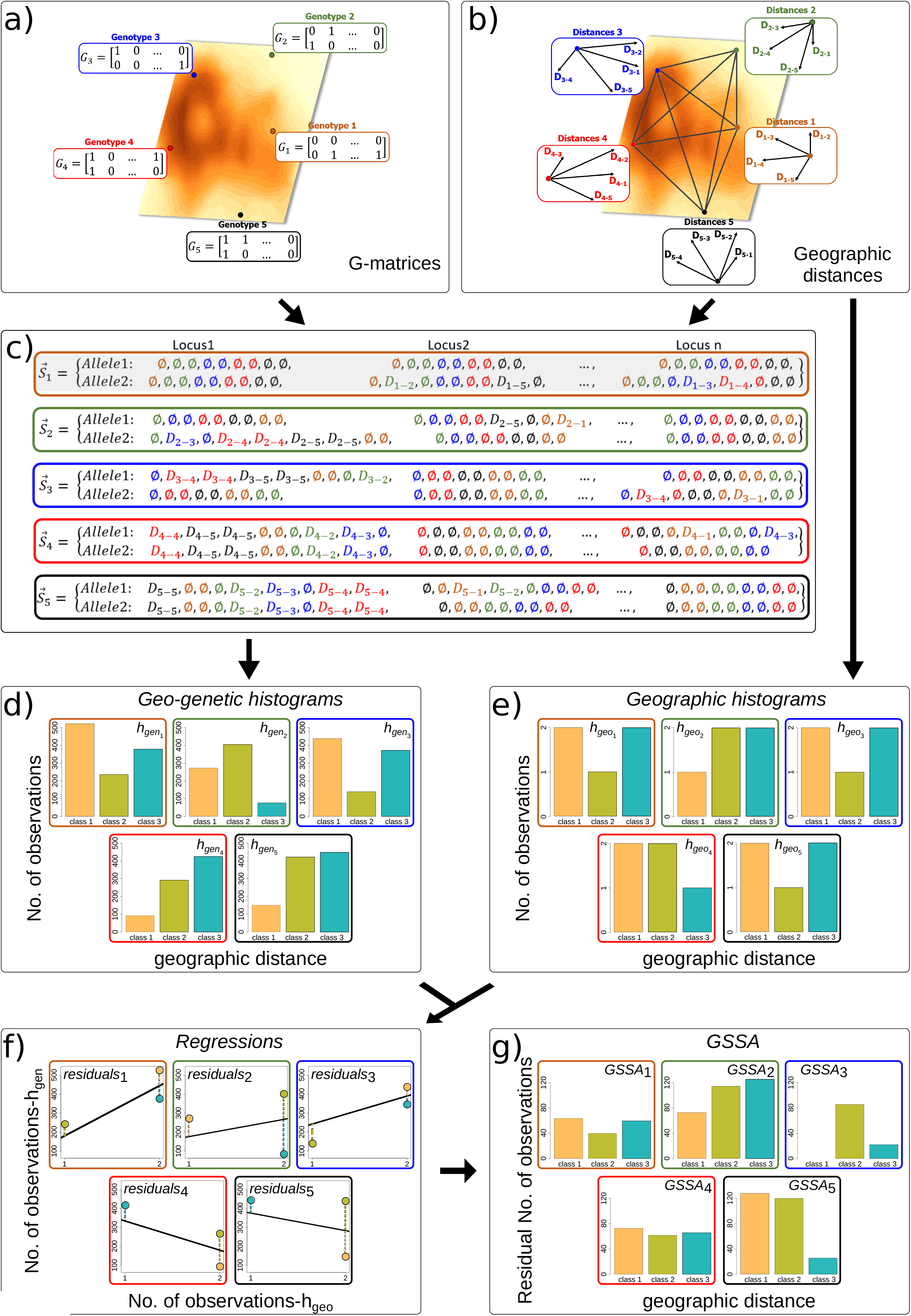
Schematic cartoon describing the construction of the geographic spectrum of shared alleles (GSSA) across five diploid genotypes that are each sampled from a unique location. Location-specific genotype matrices (*G_i_*), which capture the presence/absence of minor alleles at each location, locus and DNA strand (**panel a**), and cartoon representation of location-specific vectors (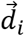), which capture the geographic distances between each focal location and all of the other locations in the sample (**panel b**). Based on the G_i_ matrices and 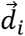 distance vectors, corresponding vectors (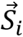) are constructed from the aggregated relative spatial distribution of minor alleles for each location (**panel c**). To facilitate interpretation, the color of each 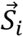 element follows the location colors in panel a, and indicates their derivation with respect to each one of the sampled locations. The 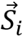 vectors are then converted into corresponding geo-genetic histograms (*h_gen_i__*), by applying a common binning scheme based on Sturge’s (1926) equation (**panel d**). For each location, a corresponding geographic histogram (*h_geo_i__*) is constructed, using the same binning scheme as before, from the geographic-distances separating the location from all other sampled locations (**panel e**). The number of observations for corresponding distance classes in both histograms,*h_gen_i__* and, are then regressed against each other for each location (**panel f**). From these regressions, the GSSA vector for each individual is built by taking the absolute value of the regression residuals for each histogram’s distance class (**panel g**).

The information captured in the GSSA helps identify the relative position of each sampling location with respect to the expansion sourcefront axis, because the similarity in genetic constitution between locations depends on the extent of shared history under an expansion history (Peter & Slatkin 2015 Bradburd *et al.* 2016). Because the amount of genetic drift increases with distance to the source of an expansion (Slatkin & Excoffier 2012 Peter & Slatkin 2013), and because colonization through different geographic paths should lead to allelic segregation (Hallatschek *et al.* 2007 Knowles & Alvarado-Serrano 2010 François *et al.* 2010), the amount of shared genetic variants can potentially further inform about the direction of the expansion. As such, if this information is captured by our summary statistic vector, it could be useful in a simulationbased statistical method—such as approximate Bayesian computation or supervised machine learning—for testing alternative expansion histories (Pudlo *et al.* 2015 Joseph *et al.* 2016 Schrider & Kern 2018).

### GSSA overview

The calculation of the set of GSSAs, one per unique sampled georeferenced location, requires two sets of data: i) the geographic distances between all sampled locations, and ii) a record of the counts of minor allele copies of each SNP locus at each sampled location (Fig. 1a). Here, we restrict our discussion to geographic distance between samples, but note that users could use an alternative distance metric, such as riverdistance, when appropriate. Below we verbally summarized the steps involved in the calculation of the GSSA for one sampled location (a formal mathematical description is presented in the next section).

First, using the entire set of geographic distances between all sampled locations, the Sturges (1926) equation is deployed to identify the optimal number of geographic distance classes and breakpoints for the construction of all of the location-specific GSSAs. This Sturges binning scheme is then applied to the set of geographic distances associated with the *i*th focal sampled location, such that there will be a specific histogram (*h_geo_i__*) for each of the locations (Fig. 1e). Second, a matrix of minor allele’s presence/absence (G_i_) is constructed for the *i*th focal sampled location (Fig. 1a). Third, a vector (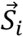) is constructed for the *i*th focal sampled location from the geographic distribution of minor alleles. Specifically, each location-specific 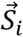 vector lists the geographic distances between each minor SNP allele present at the focal sampling location and all other copies of the same minor allele in the entire sample (Fig. 1c). Fourth, this location-specific vector is then converted into a corresponding location-specific histogram (*h_gen_i__*) using the Sturges’ (1927) binning scheme previously defined (Fig. 1d). Finally, to make our statistic independent of the specific sampling scheme (i.e., sampling locations and their influence on the binning scheme), the location-specific histogram constructed from the vector of distances between minor alleles (*h_gen_i__*) is regressed against the values from the corresponding “geography-only” histogram (*h_geo_i__* i.e., the “null” distribution of geographic distances). The residuals from this regression are finally used to build the location-specific histogram for the *i*th focal location (GSSA_i_).

### GSSA calculation

Formally, the two sets of data extracted from the series of georeferenced sampled genotypes can be defined as two matrices. With one individual per sampled location, *I* locations, *M* sets of chromosomes (i.e., ploidy), and *L* loci (i.e., SNPs), the arrangement of minor alleles in the entire sample relative to each location can be characterized by a *M* x *L* genotype matrix, *G_i_*, where each element *G_i_m,l__* takes the value of 0 or 1, depending on whether a copy of the minor allele is present (1) or not (0) at each individual’s SNP locus and DNA strand (note that phasing is not necessary because the method treats each locus independently Fig. 1a). In turn, the set of Euclidean or effective (Shirk & Cushman 2014 Davis *et al.* 2018) geographic distances between all locations can be characterized by an *I* x *I* square distance matrix, *D*, where D_i,i_ = 0 and *D_i,j_* = *D_j,i_*. From this latter matrix, a set of location-specific geographic-distances vectors, 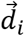, can be obtained by subsetting the D matrix so that 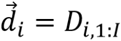 (Fig. 1b). Altogether, these data are used to construct a GSSA summary statistic vector for each sampled location following five steps:

*Step 1.* First Sturges’ (Sturges 1926) equation is applied to the set of pairwise geographic distances between all localities contained in matrix *D* to identify an optimal series of consecutive distance classes that are later used for histogram binning. We chose Sturges’ (1926) equation for its simplicity and common use in biological research (RamírezGarcıa *et al.* 1998 Fagua & Gonzalez 2007 Pires *et al.* 2016 Cardozo *et al.* 2018). Although alternative binning schemes would in theory be possible, an exploration of a coarser binning scheme (Scott 1979) and the more granular Rice rule (Jones *et al.* 2001--) showed that although Sturge’s scheme resulted in relatively better performance with regards to the dynamics of the GSSA and identification of expansion colonization history, accuracy was good across all three binning schemes (Supp. Fig. 1).

*Step 2.* Each genotype matrix *G_i_* is used to construct a location-specific vector, 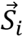, which summarizes the aggregated relative spatial distribution of minor alleles at location *i* (Fig. 1c). The elements of this vector are defined as:

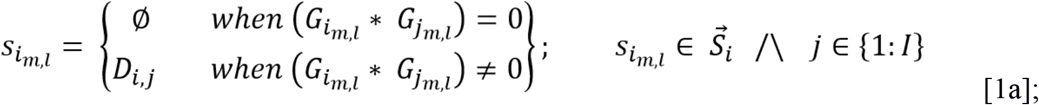

where Ø = null, *G_i_m,l__* = element of individual *i*’s genotype matrix that denotes the presence or absence of a minor allele at locus *l*, strand *m*, and *D_i,j_* = geographic distance between localities *i* and *j.* Elements in the 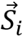 vector with a value of zero correspond to instances in which the same minor allele at a SNP locus is homozygous within an individual or present in more than one individual sampled from the same locality because the geographic distance between these allele copies would be zero (D_i,i_ = 0). On the other hand, nonzero elements correspond to the geographic distances between individuals from different localities sharing the same minor allele at a SNP locus (Fig. 1c). Effectively, 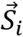 can also be derived from the frequency of the minor allele at each locus and location by aggregating over DNA strands so that the set of 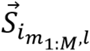 elements of this vector are defined as:

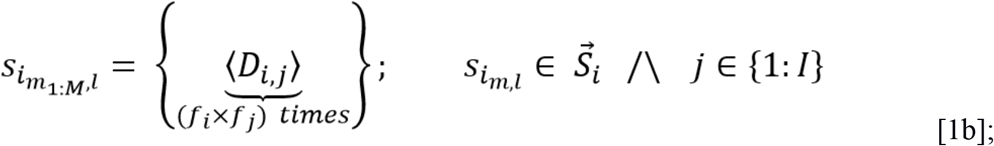

where *f_i_* is the frequency of copies of the minor allele at location *i*, locus *l* and *D_i,j_* as in eq. [1a].

*Step 3.* After removing all null elements (Ø) from vector 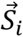 (Fig. 1c), which number depends on the number of loci at which each locality does not contain a minor allele copy (Fig. 1a), the size of each 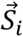 varies among locations. Therefore, it is necessary to compress each 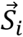 into a vector with an equal number of elements for all individual sampled locations. To do this, each vector 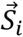 (Fig.1c) is converted into a histogram by calculating the frequency of 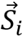 elements that fall within each of the distance classes previously determined in *Step* 1. The resulting location-specific geo-genetic histogram (*h_gen_i__*) summarizes the relative spatial distribution of minor alleles copies across loci present at each (*i*^th^) sampling location (Fig. 1d).

*Step 4.* To correct for the “null” geographic expectation introduced by the specific geographic position of each sample, each location-specific vector of geographicdistances, 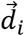, is first converted into a histogram by again estimating the frequency of observations that fall within each of the same set of distance classes previously used to create the geo-genetic histograms. This generates a unique geographic histogram (*h_geo_i__*) for each (*i*^th^) location that is analogous to the corresponding geo-genetic histograms (*h_gen_i__*) (Fig. 1e), yet that only containing geographic information.

*Step 5.* Finally, the elements of each geo-genetic histogram (*h_gen_i__*) are regressed against the corresponding elements of the geographicdistance histogram (*h_geo_i__*) (Fig. 1f), as defined here:

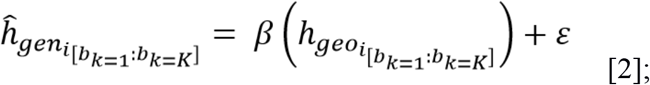

where *h_gen_i__* = geo-genetic histogram for location *i*, *h_geo_i__* = geographic histogram for location *i*, [*b_k_*] = histogram’s distance class, *b*, ranging from k = 1 to k = K (i.e., the maximum number of bins as determined by Sturges’ equation), β = simple regression coefficient, and ε = error term.

The vector of absolute residuals resulting from each regression (eq. [2]) constitutes the location-specific spatial summary statistic vector we named the GSSA and it is defined for each location *i* as the absolute difference between the location-specific regression-predicted values (eq. [2]) and the location-specific geo-genetic histogram values:

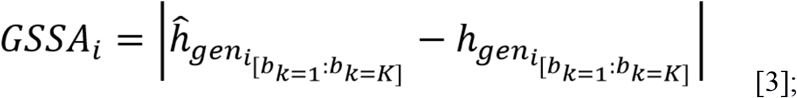

with *ĥ_gen_i__*, *h_gen_i__*, and [*b_k_*] defined as in eq. [2].

### GSSA Implementation

All five steps described above can be implemented from a list of genotypes and a list of individuals’ sampling coordinates, or optionally a set of distances between genotypes in case users choose to use effective distances instead (Shirk & Cushman 2014 Davis *et al.* 2018). The aggregated relative spatial distribution of minor alleles vector for each locality,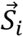, can be obtained from the multisite frequency spectrum (multiSFS), which summarizes the frequency of shared allele variants across populations, and the set of distances among all sampled locations. This multiSFS can be calculated from the list of genotypes in available programs such as ***δ****a****δ****i*, whereby the multiSFS should be polarized based on the minor allele where each sampled georeferenced location is considered a “population” in the multiSFS (Gutenkunst *et al.* 2009). Scripts to estimate the multiSFS along the rest of necessary steps to calculate the GSSAs for each location are available at https://bitbucket.org/diegofalvarados/gssa-v0.0/src. If multiple individuals per location are available, the method can be adapted by aligning all genotypes per location into a single “polyploid individual” so that individuals’ genotypes are treated as different DNA strands. Similarly, polyploid individuals are likewise accommodated as the method treats each SNP locus independently and thus, no phasing is necessary.

### Raggedness index

A useful property of the GSSA is that its shape can potentially carry information about a location’s relative age of colonization and its historical connectivity with other populations given an expansion history. To explore and summarize the overall behavior of the GSSA with respect to expansion sources, we use Harpending’s raggedness index (Harpending 1994). Harpending’s raggedness index (RI) was originally introduced to quantify the shape of the histogram obtained from the average pairwise genetic differences amongst individuals in the context of inferring the history of population size change due to the predictive relationships between the shape and the timing and magnitude of size change. Here we repurpose it to condense the GSSA elements into a single variable that is correlated with the shape of the GSSA and thus, summarizes the behavior of the GSSA and its spatial and temporal relationship with the expansion source. In our calculation of each localityspecific RI, we disregard all distance classes within each GSSA_i_that are equivalent to the distance classes in the corresponding *h_geo_i__* that have a value of zero (which arise for distance classes not involving the focal locality). We then correct the RI for the number of comparisons used for its calculation:

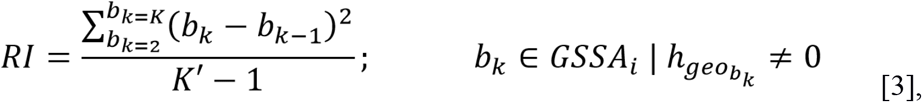

where *b_k_* indicates a distance class and *K*′ the total number of distance classes used in the calculation.

Locations further away from the serialexpansion source are expected to share more genetic variants with nearby locations that were colonized through the same expansion path (Knowles & Alvarado-Serrano 2010), and hence should tend to present a leftskewed and more ragged GSSA (Supp. Fig. 2a). This is because, in this case, short distance bins should be overly represented with a marked drop at intermediate bins, thereby inflating the RI from the “neighborhood effect” resulting from allele surfing (Hallatschek & Nelson 2008 François *et al.* 2010). This process can cause genetic variants at the expansion front to increase in frequency within constrained areas of derived populations (Excoffier *et al.* 2008). In contrast, locations closer to the source should tend to have a more uniform and nonskewed GSSA histogram because these locations are expected to retain the shared genetic variants present at different frequencies in other locations in the sample. In this latter scenario (Supp. Fig. 2b), most variants come from standing genetic variation in the source population due to the comparatively smaller amount of drift experienced by these closetosource populations (Excoffier *et al.* 2008 Peter & Slatkin 2013).

### Simulation experiments

To evaluate the behavior of the GSSA, we simulated 5,000 unlinked SNPs sampled from 10 spatially separated individuals under different serial range expansion histories, where 99 demes are colonized by a single source deme. Spatially implicit simulations were conducted using fastsimcoal v2.5.2 (Excoffier & Foll 2011 Excoffier *et al.* 2013), with each simulation starting with a source deme at either of four different starting locations (Fig. 2). At each sequential expansion step going forward in time, we set a proportion of individuals within each deme (*f*) to move out from the demes they occupied into an immediate neighboring deme, excluding diagonal movements (i.e., 4-neighbors). We set newly colonized demes to then grow to a size *N_K_* (based on the carrying capacity parameter), in *τ_r_* generations. After *τ_c_* generations, we set the next set of demes to be colonized from previously colonized populations, and cycles of serial colonization proceed until the last of 99 demes is colonized. We set the simulations to then run for τ additional generations after the last colonized deme has reached *N_K_*. Through the entire simulation, we allowed colonized demes to exchange individuals with neighboring colonized demes (4-neighbors) according to a migration parameter (*m*) that reflects the per individual per generation probability of migration. After all available demes are colonized, we let the simulations continue under a steppingstone model (Kimura & Weiss 1964) for a number of generations determined by parameter (*τ*), which signifies the time after initial expansion has ended. Table 1 lists all parameters controlling the simulations.

**Fig. 2.**
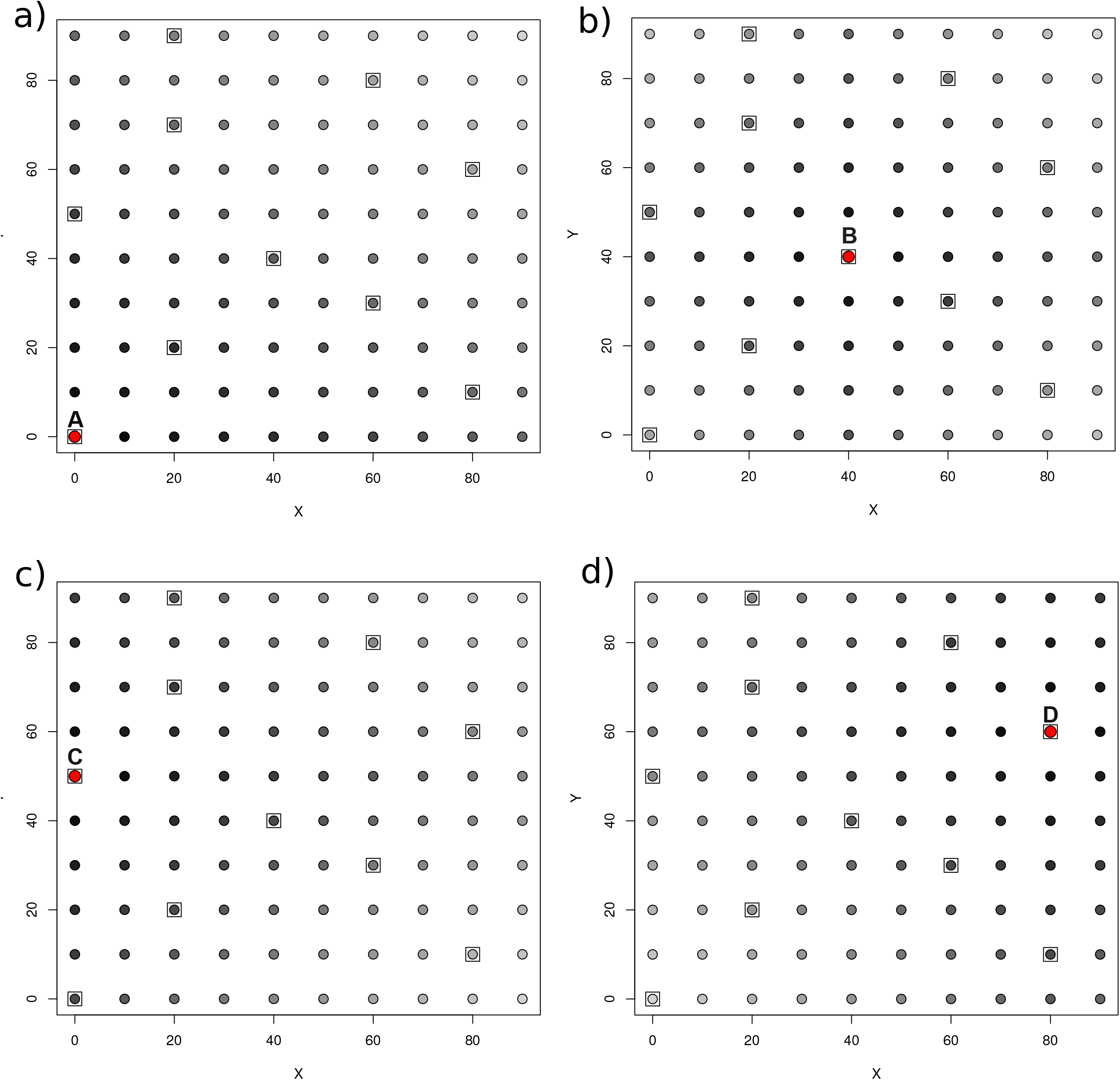
Schematic of the sequential range expansion models used to quantify the spatial association of the GSSA and expansion source location. Each panel correspond to simulations started from a different source (marked in red and denoted by letter). Demes are shaded according to their colonization time, with darker colors corresponding to earlier colonization times. Sampled demes are surrounded by a square.

**Table 1.** Parameters used in simulations. Parameter values in bold were used for all simulation experiments except those in which the parameter is being tested.

| Parameter | Description | Values tested | Unit |
| --- | --- | --- | --- |
| $\tau$ | Time allowed for the simulation to continue after last deme has been colonized and reached its carrying capacity | **0.0**<br>0.5<br>1.0 | Coalescent units |
| $f$ | Proportion of individuals of a source deme that settle each uncolonized deme | **0.002**<br>0.01<br>0.1 | Proportion of $N_K$ |
| $\tau_r$ | Time allowed for demes to reach $N_K$ after being colonized | **0.01**<br>0.05<br>0.10 | Coalescent units |
| $m$ | Pairwise probability of migration per individual per generation between neighboring colonized demes | **0.000**<br>0.001<br>0.010 | Proportion |
| $N_K$ | Effective population size of deme at carrying capacity | **1000** | Number of individual |
| $\tau_c$ | Time lag between source deme reaching $N_K$ and colonizing a new deme | **0.01** | Coalescent units |

The particular effect of each parameter on the ability of the GSSA vector (summarized with the RI) to locate the expansion source was quantified independently. For that, we kept all parameters—except the one of interest of interest—fixed across simulations (Table 1). For each scenario/parameter combination, we ran 1,000 simulations for a total of 48,000 simulations (1 scenario x 4 possible sources x 4 parameters x 3 parameter values x 1,000 simulations). After each simulation was completed (as defined by *τ*), we gathered a dataset of 5,000 homologous biallelic SNPs from one individual from each of the 10 locations that were randomly selected before the simulations, including the source location. We then used our custom python pipeline to calculate a GSSA (eqs. [2, 3]) and associated RI (eq. [4]) for each of the 10 sampled locations. The source of the expansion was identified based on the distribution of magnitudes of the RI across sampling locations, with the smallest RI value assumed to correspond to the expansion source. Accuracy was measured as the proportion of simulations that correctly identified the source. To capture the empirical reality that the actual expansion source may not have been sampled, we repeated our inferences while excluding the source location from our sample (Fig. S2). In this latter case, accuracy was measured as the proportion of simulations that correctly identified the most proximate location to the source location among those sampled (Supp. Fig. 3). In the case of the simulations that include the source, the root mean squared error (RMSE) was also calculated. Additionally, we visualized the magnitude of change in the RI of each sampled location as a function of when each location was colonized.

As a point of comparison, we also used Peter and Slatkin’s (2013) approach to identify the expansion source on the same simulated datasets. Because Peter and Slatkin’s approach make use of interpolation, which reduces the chances of estimating the precise coordinates of the simulated source, we scored a prediction as accurate whenever Peter and Slatkin’s directionality index was able to locate the source within the perimeter defined by the 8neighbor demes around the actual source.

### Empirical Application

To illustrate and demonstrate the utility of our approach, we applied it to two taxa of the holarctic butterfly genus *Lycaeides* that are distributed in western North America (Gompert *et al.* 2014b). These butterfly taxa are hostspecialists and relatively poor dispersers, with distributions that are tightly linked to their host plants (Forister *et al.* 2011 Gompert *et al.* 2014a). Their respective geographical ranges were likely impacted by Pleistocene climatic events (Thompson *et al.* 1993 Thompson & Anderson 2000 Pierce *et al.* 2004), which presumably involved range expansions from refugia since the Last Glacial Maximum (LGM). In particular, we focused on two highelevation undescribed taxa (Alpine and Jackson *Lycaeides* Gompert et al. 2014). These data are wellsuited for demonstrating our approach because they have a relatively wellcharacterized evolutionary history and spatial distribution (Gompert *et al.* 2006 Nice *et al.* 2013), as well as a thorough, spatially widespread genetic sampling (Gompert et al. 2014) in the western mountains of North America (Alpine *Lycaeides*: 10 localities and 8097 independent homologous SNPs Jackson *Lycaeides*: 11 localities and 9074 independent homologous SNPs).

To compare our geographic predictions of source locality with the plausible geographic origin of each taxon’s putative range expansion after the LGM, we used ecological niche models (ENMs), projected onto past climates, as independent hypotheses of colonization history under the assumption of Grinnellian niche conservatism. Briefly, we generated an ENM for each taxon in the R package dismo (Hijmans 2012) using a maximum entropy approach (Phillips *et al.* 2006) and 19 interpolated climatic surfaces at 30 arc-seconds resolution that summarize global patterns of temperature, precipitation, and seasonality (Hijmans 2012). We only included localities for which genetic data are available (Table 1, Gompert et al. 2014) given the pending taxonomic status of these taxa, which can lead to misidentification issues in available locality databases. To reduce possible sampling bias, for each taxon we filtered the localities based on a minimum distance required between localities. The exact distance used for each species was determined by a variogram approach that establishes the distance over which spatial autocorrelation in environmental conditions is minimal (Brown *et al.* 2016). In addition, to maximize the fit of the model and to avoid unnecessary complexity, we tuned our models using the R package ENMeval (Muscarella *et al.* 2014). The tuning procedure consisted of assessing optimal model parameters based on jackknife crossvalidation of the samples (Radosavljevic & Anderson 2014 Muscarella *et al.* 2014). We chose a jackknife approach given the small sample size of the filtered collection points (Galante *et al.* 2018). Model fit was evaluated under different combinations of model features and regularization multipliers (Table S1) using three sequential criteria: omission rate, AUC, and model feature class complexity. Tuning results are summarized in Supp. Table 1. We then hindcasted this ENM to the LGM using climatic CCSM (Community Climate System Model) estimations derived from the Paleoclimate Modelling Intercomparison Project Phase II (Braconnot *et al.* 2007). Finally, following Knowles and Alvarado-Serrano (2010), we identified the area(s) of contiguous high predicted suitability (i.e., refugial distribution) as potential expansion source(s), using as an arbitrary cutoff the top 10 ^th^ percentile of the predicted suitability scores. We then contrasted these hypothesized sources with those inferred by our approach.

## Results

### Simulation experiments

Our simulation study showed that the magnitude of the RI calculated from the elements of our GSSA vector was highly correlated with i) the distance from the location of the expansion source, and ii) how long, after the expansion started, a location was colonized (Fig. 3). The metric was able to accurately identify the geographical source of expansion given that one of the sampling localities was the source of expansion, or the first location colonized among those sampled (Fig. S3). Still, the degree of accuracy somewhat depended on the location of the source deme such that lower accuracy occurred when the source was located in a peripheral area (Fig. 4). Still, incorrect inferences tended to identify sampling localities proximate to the actual expansion locality, regardless of the location of the expansion source (Fig. 5).

**Fig. 3.**
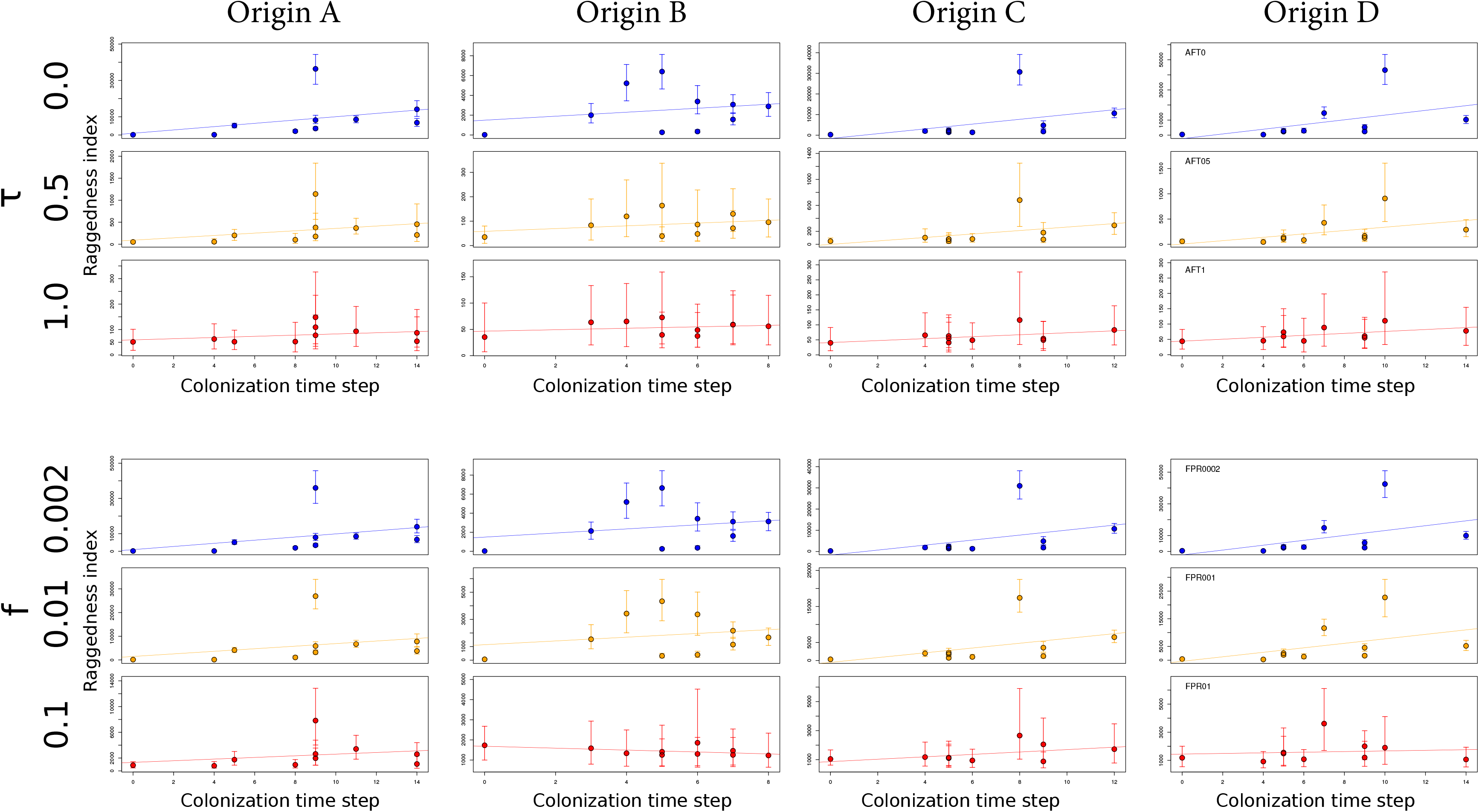
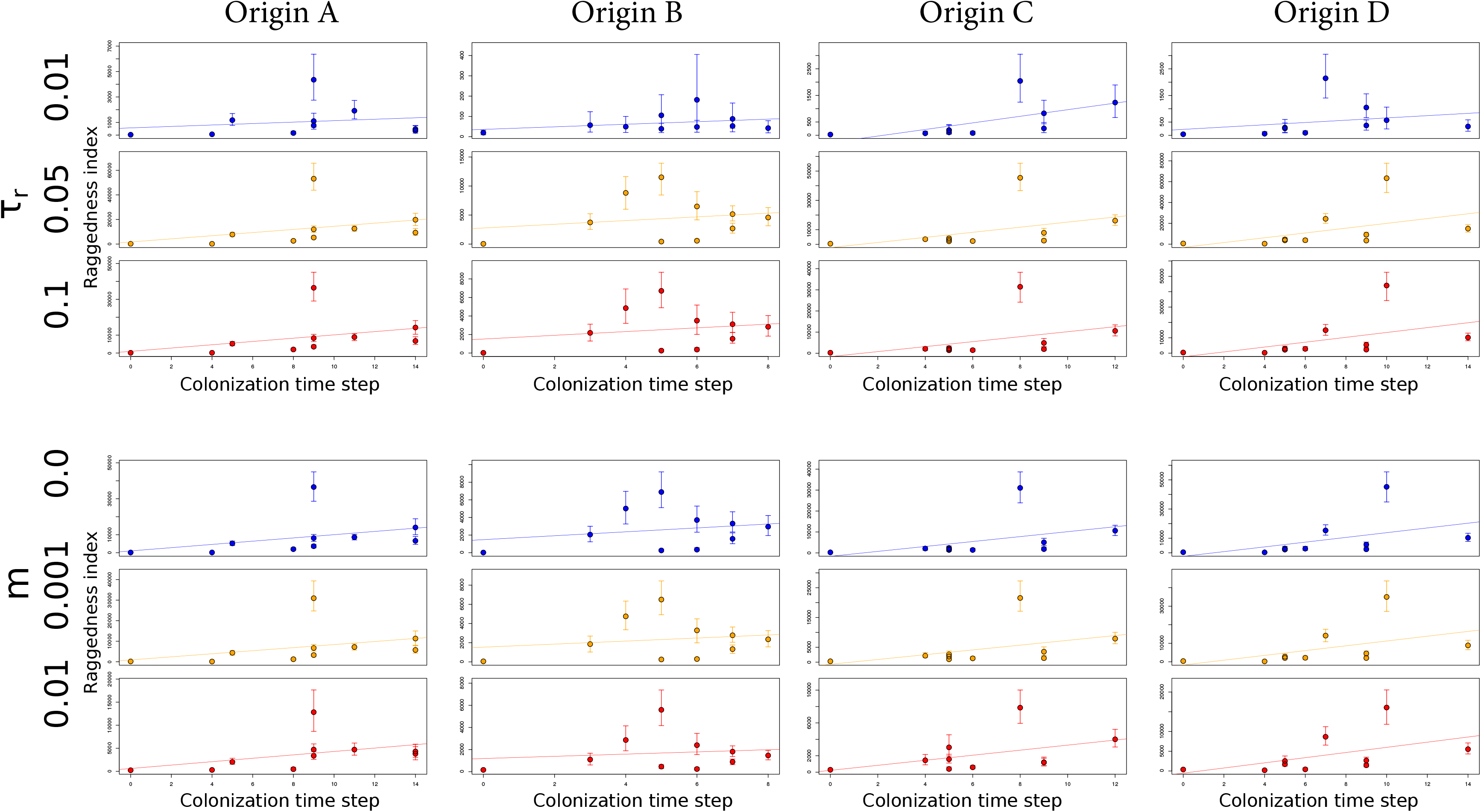
Association between time at which each sampled locality is colonized and the raggedness index calculated from the GSSA at each of these localities under all four simulated scenarios. Colonization time step denotes the time at which a locality was colonized (with the origin always being zero). The impact of each parameter on the ability to recover the simulated colonization history is evidenced by changes in the slope of the association. Different colors are used for each parameter value for increased visibility.

**Fig. 4.**
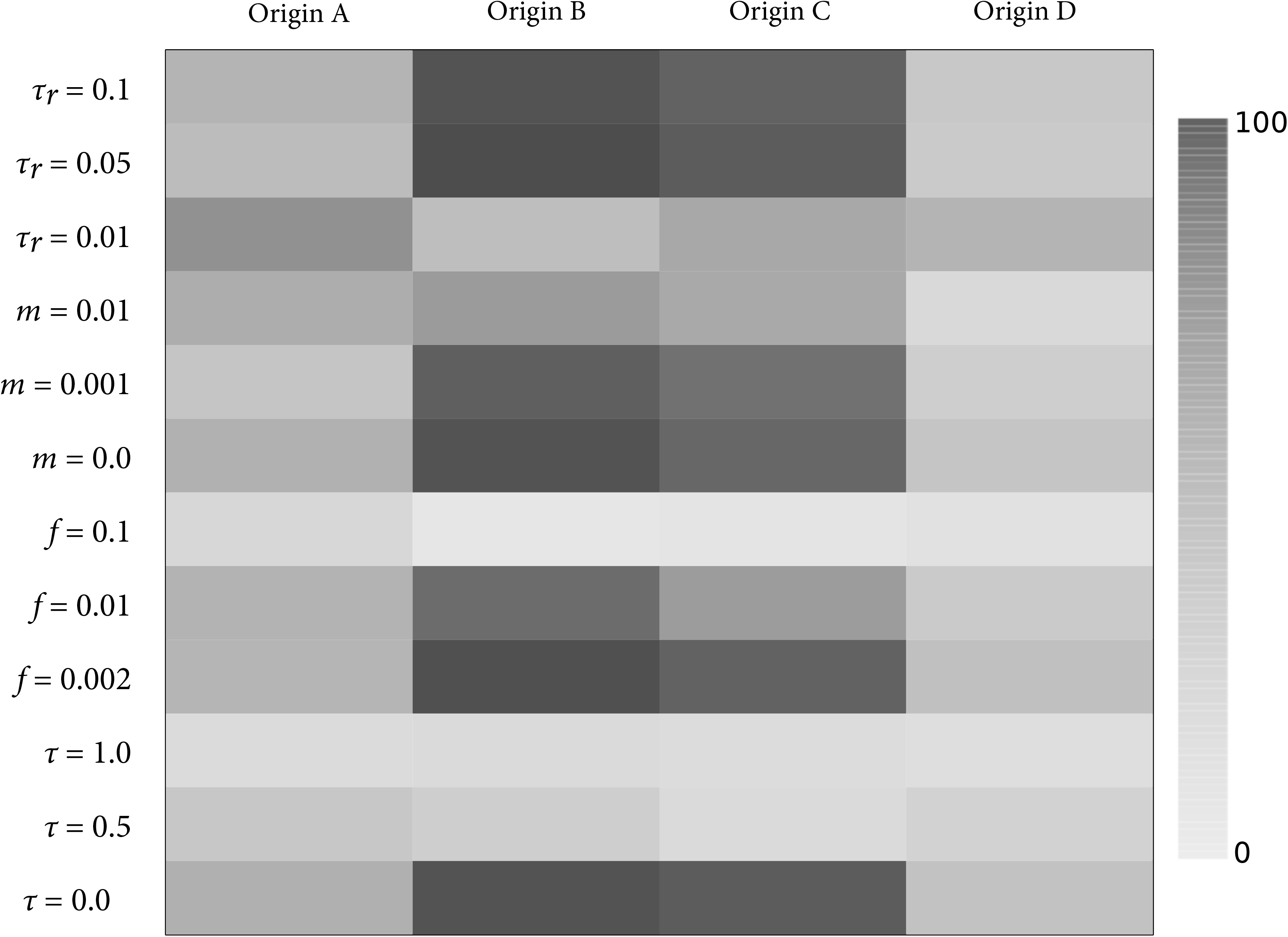
Probability across 1000 simulation replicates of the the raggedness index calculated from the GSSA vector for identifying the source deme given the four serial range expansion scenarios and model parameter values. Higher probabilities of correct source identifications are a function of darker shadings. See Table 1 for model parameter definitions.

**Fig. 5.**
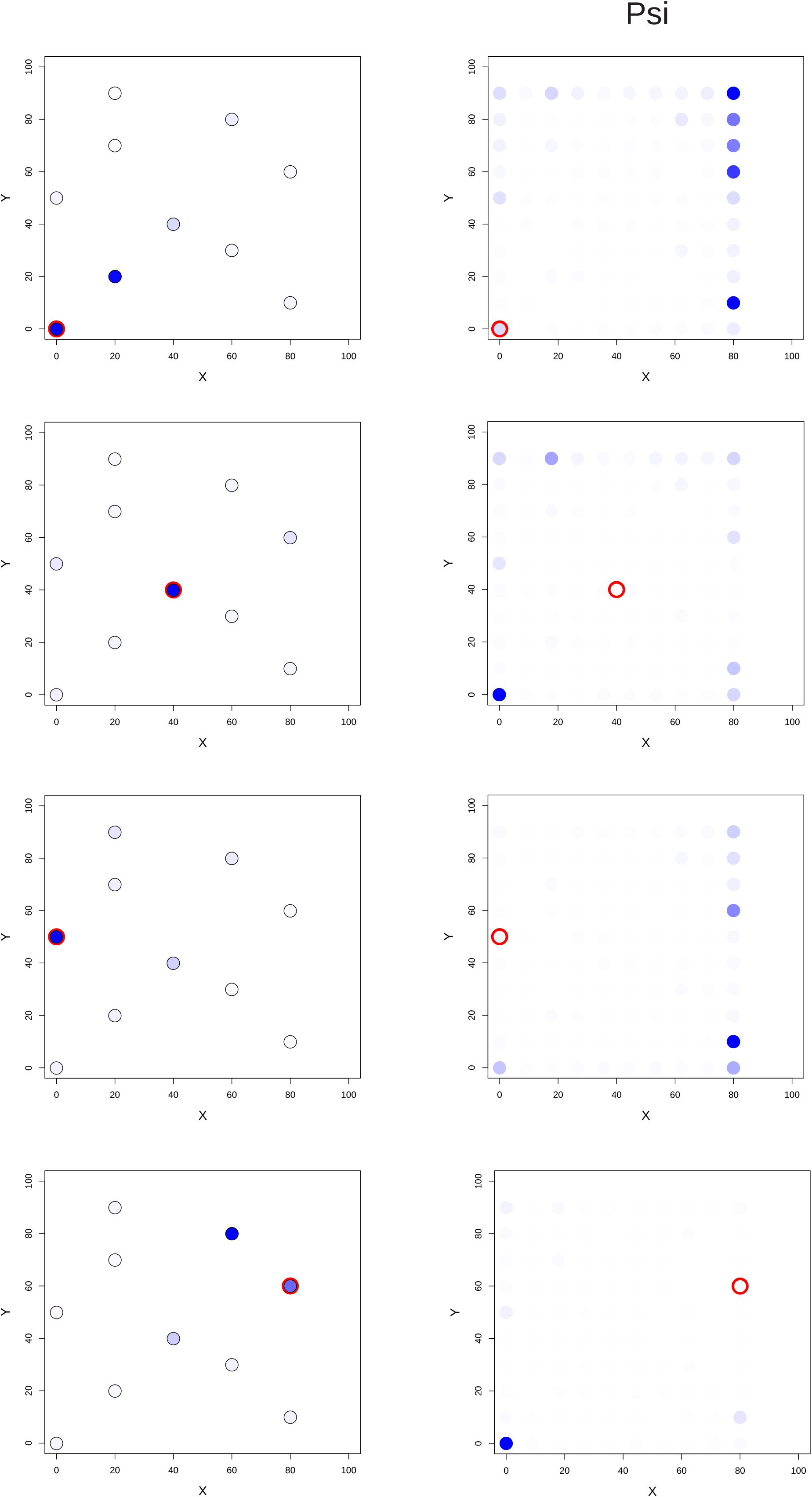
Geographic position of the suggested sources of expansion according to the raggedness index calculated from the GSSA vector (left column) and directionality index (right column). A red circle denotes the true source used for each simulation scenario, whereas black circles indicate the locations sampled (shown only for the GSI-RI approach). Results presented are aggregated across 1000 simulations, with the relative frequency that each position was identified as the source of expansion indicated by the color hue (darker colors corresponding to greater frequencies). Note that in the vast majority of instances, the GSS-ARI approach either correctly infers the source or identifies a close neighbor location.

By summarizing the GSSA with Harpending’s (1994) RI, we were able to correctly identify the source population up to 70% of the time, with RMSE values that approached zero (Fig. 6). However, this accuracy declined with larger numbers of colonists (*f*) and time since all demes have been colonized (*τ*). Likewise, under an IBD model that arose when enough time had accrued for the signature of the expansion to have significantly diminished (Fig. 6), our metric was unable to accurately identify the source population. Our approach did strongly outperform Peter and Slatkin’s directionality index (ψ) by a margin of up to 60%, except in those aforementioned cases such as a high number of individuals colonizing demes (Fig. 6).

**Fig. 6.**
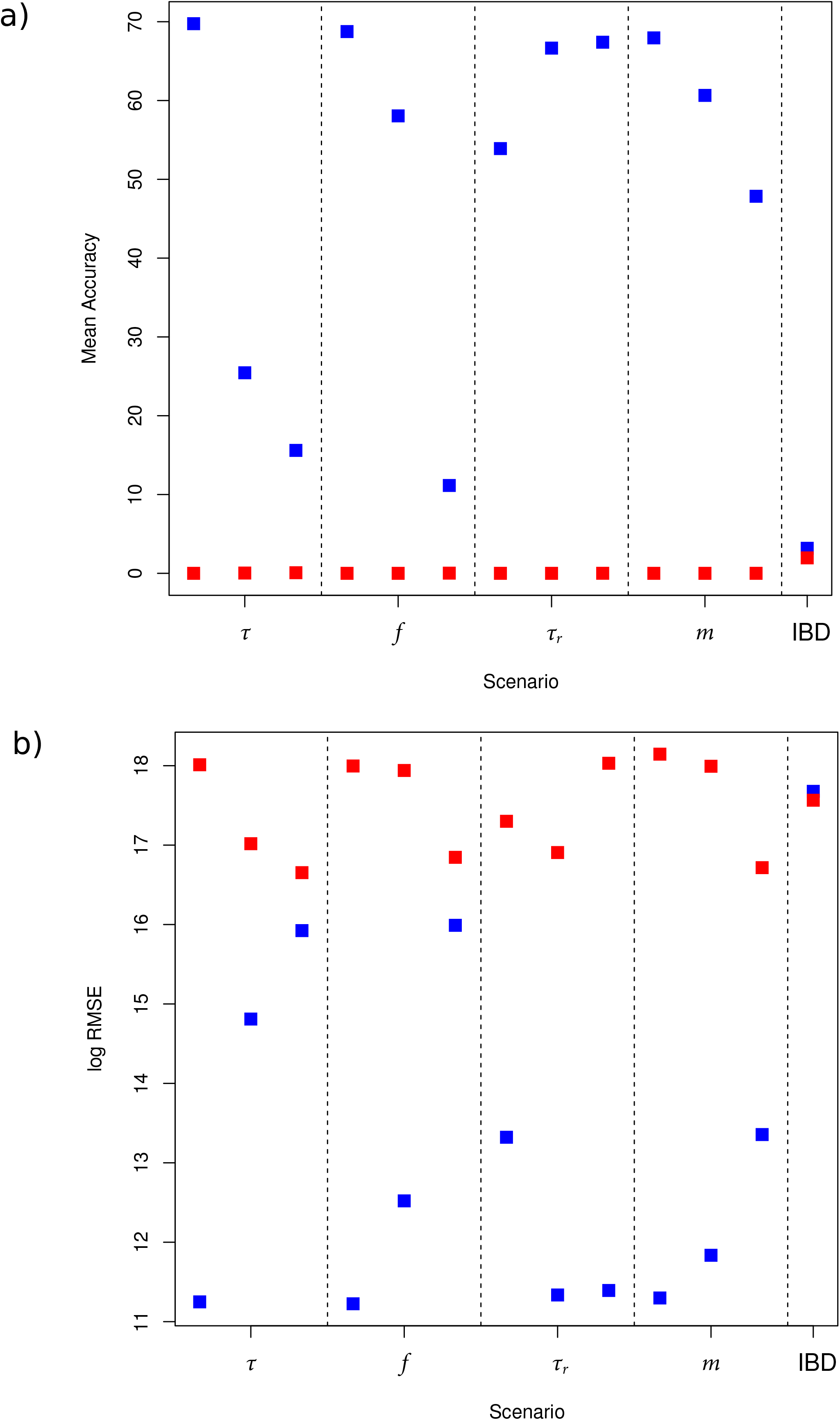
Comparative accuracies of the raggedness index calculated from the GSSA and directionality indexin identifying the source deme for serial range expansions. The mean accuracy (a) per model parameter tested and corresponding root mean square error (RMSE) (b) obtained under both methods (GSSA-RI in blue, directionality index in red) is shown under different model parameter values.

### Empirical application

The application of our coupled GSSARI approach to the two *Lycaeides* datasets suggests an expansion spatial dynamics that is well in line with species ranges at the LGM according to our ENM hindcasts (Fig. 7). For the Alpine *Lycaeides* distributed in northern Rocky mountain areas, the predicted source coincided with the southernmost samples, which are located in the largest LGM refugium predicted by the ENM hindcasts (Figure 7A). Although the hindcasted range at the LGM climate for this species covers a wide area that encompasses all of the sampled locations, the sampling location predicted by the GSSA coincides with an area of highest suitability. The RI calculated on the GSSA for Jackson *Lycaeides* samples suggested a source also within the ENM hindcast (which was more heterogeneous in comparison with Alpine *Lycaeides* hindcast), and close to one potential refugium (i.e., continuous area of high suitability) (Fig. 7B).

**Fig. 7.**
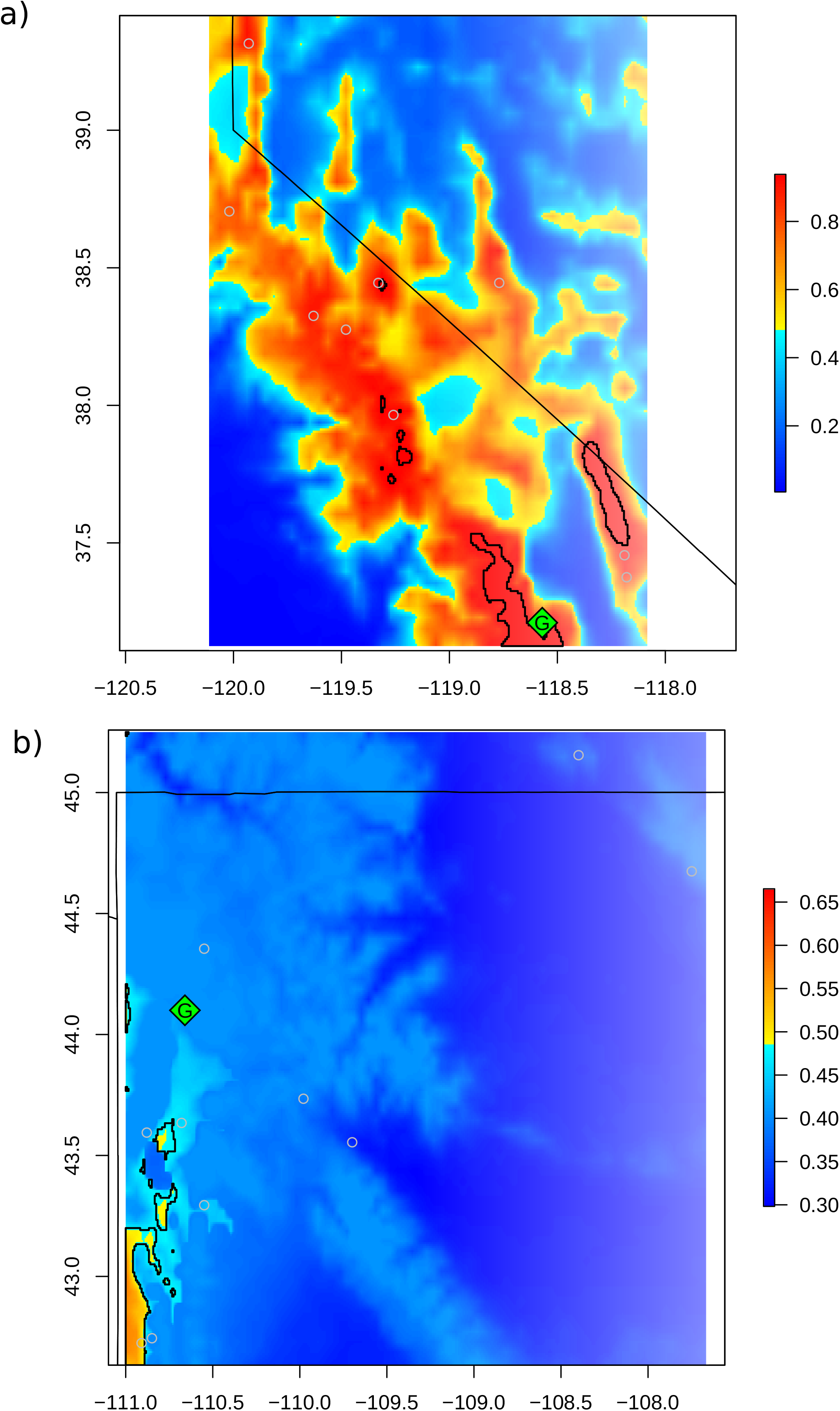
Comparison of LGM hindcasts of range distributions obtained with Ecological Niche Models (ENMs). The grey circles indicate the sampling localities on which the inferences are based. The suggested sources of expansion according to the GSSA-RI approach are indicated by a green rhombus with a “G” on it. Warmer colors indicate greater potential suitability as estimated by the ENM. a) Model and inferences for Alpine *Lycaeides* b) model and inference for Jackson *Lycaeides*. Refugia, identified as the top 10th percentile most suitable areas, are denoted by black polygons.

## Discussion

We introduce a new summary statistic vector, the geographic spectrum of shared alleles (GSSA), which makes joint use of geographic and genetic information. We demonstrate that it can be used to help infer the geographic dynamics of a range expansion. We use Harpending’s (1994) RI calculated on the GSSA elements of each sampled genotype to summarize spatial gradients in the distribution of minor alleles that are indicative of the relative position of each genotype along an expansion axis. However, we envision the elements of the GSSA vector may be directly incorporated within a supervised machine learning or ABC inferential framework to obtain parameter estimates associated with source locality likelihoods, and to test competing expansion hypotheses (He *et al.* 2017b Fraimout *et al.* 2017 Schrider & Kern 2018).

However, there are some important assumptions to first examine before deployment of the GSSA. First, as with many other population genomic approaches, it should be used after population structure is explored (Patterson *et al.* 2006 Frichot & François 2015 Petkova *et al.* 2016). Specifically, the GSSA is applied to groups of samples suggested to derive from single sources by various other exploratory methods applied cautiously (House & Hahn 2017 Elleouet & Aitken 2018). For example, as late Pleistocene range expansions and recent species invasions can both have multiple origins (Miraldo *et al.* 2011 RuizCooley *et al.* 2013 Cristescu 2015), care needs to be taken to detect the possibility of admixture from multiple origins *a priori*. In this case, if exploratory methods suggest a species is heavily structured, with a history of expansions from more than one refugium, researchers could apply the GSSA to each subsample independently. This may, however, prove a challenging task in certain circumstances because smaller sample sizes (resulting from splitting up the range) may lead to reduced accuracy. Difficulties may also arise if more than two sources are close together geographically, multiple expansions occurred, or if gene flow directions are biased given complex heterogeneous geography (Branco *et al.* 2018 Lundgren & Ralph 2018). However, one strength of the approach is that the assumption of panmixia within each sub-sample can be relaxed, as our simulated conditions contain stepping stone relationships among demes such that some isolationbydistance relationships within genetic clusters is accommodated, as long as there is a single source.

A second assumption underlying our metric is that a single spatial expansion had actually occurred within an identifiable timeframe. As demonstrated by our simulations with the reduced accuracy as the time since all demes have been colonized (τ) increases, eventually reaching the stationarity conditions of isolationbydistance, the historical signal of expansion will be erased over long intervals. It is therefore recommended that spatial expansion be tested before attempting to infer the geographic dynamics of an expansion (Excoffier 2004 Wegmann *et al.* 2006 Elleouet & Aitken 2018).

Key advantages of the GSSA are that no outgroup is required, and that it can be deployed on sparse datasets that are often collected from a variety of nonmodel species. Notably, correlation to the source of expansion was in general robust to different acrosspopulation migration rates (*m*) and deme growth rates—as conditioned by the time allowed for demes to reach their carrying capacity (*τ_r_*). However, accuracy did somewhat decrease with greater migration levels, as one would expect. Also expectedly, the accuracy in identifying the source location was greatly diminished by larger founder sizes (*f*), and the amount of time passed after colonization ended (*τ*). In the former case, larger *f* will deflate the founder effect, whereas in the latter case the signal of the founder effect will become erased over time (Nei *et al.* 1975).

Regardless of parameter setting, the GSSA approach consistently outperformed the directionality index of Peter and Slatkin’s (2013) in all simulations. While such a finding is not surprising given that we violated the latter method’s assumption of allelic polarity and used smaller sample sizes than recommended, it highlights the gap filled by our new statistic, which accommodates data typical of nonmodel organisms. As demonstrated by Peter and Slatkin’s (2013) empirical example, in which they found support for the outofAfrica hypothesis (Li *et al.* 2008 DeGiorgio *et al.* 2009) using a comprehensive empirical dataset consisting of over half a million SNPs in more than 1,500 individuals distributed in 55 human populations (Fumagalli *et al.* 2011), their statistic is a powerful tool for model organisms. Ours on the other hand, may be best suited for smaller datasets. Furthermore, it is important to note that the two metrics are not precisely inferring the same entity even if the overall objective is to uncover the spatial dynamics of an expansion. Specifically, the smallest RI of the GSSAs is assumed to be associated with the sampling location closest to the actual origin (i.e., first colonized location among those sampled). In contrast, Peter and Slatkin’s ψ aims to infer the actual geographical point of expansion, which can be located beyond the sampling localities. This methodological difference reflects our goal of limiting the amount of statistical uncertainty at the expense of decreasing potential inferential capabilities (i.e., limiting the noise introduced by spatial interpolation). Nevertheless, part of our simulation study used sampling locations that included the source, thereby making this comparison between both approaches informative.

The utility of our new statistic for nonmodel organisms is also apparent in the empirical analysis of *Lycaeides* (Fig. 7). These taxa had 8097 and 9074 independent SNPs (i.e., 1 SNP per tag for Alpine *Lycaeides* and Jackson *Lycaeides* respectively), from which we randomly selected a set of 5000 SNPs for consistency with our simulations. Our approach was able to point, as sources, geographical regions of relatively high predicted suitability according to the ENMs. For Alpine *Lycaeides*, this included a location at the southern end of the range. In contrast, for *Jackson Lycaeides*, the lowest RI suggested an area of marginal LGM suitability (albeit close to regions predicted to be suitable Fig. 7). That being said, because the inferred entity refers to the sampling location that is closest to the true origin (i.e., first colonized location among those sampled), a perfect match between the ENMs’ prediction and the GSSA inference is not expected. Together with the simulation data, these analyses demonstrate that under circumstances of more constrained information and limited sampling relative to those common to model organisms, our approach may be a useful complement to existing methods to infer the geographic history underlying range expansions (Ramachandran *et al.* 2005 François *et al.* 2010 Peter & Slatkin 2013). Our results also highlight how independent lines of evidence can be combined for more robust demographic inference. In contrast to previous attempts to reciprocally validate ENMs and genetic data without spatially explicit inference (Alvarado-Serrano & Knowles 2014), our method allows one to make more explicit comparisons between the hypothesized locations of refugia derived from ENMs and the inferred range expansion dynamics based on genetics. This capability is of particular interest when considering the uncertainty associated with hindcasting using ENMs, as these tools are capable only of generating hypotheses about potential distribution based on abiotic correlates under the assumption of niche conservatism and analogous climate—which is likely to overpredict the real former distribution of species (Soberón 2007 Peterson *et al.* 2010 Alvarado-Serrano & Knowles 2014). By combining these approaches, the confidence in these estimates increases, giving scientists the ability to refine a set of plausible historical scenarios under consideration.

### Potential Pitfalls

To help improve modelbased inferences, several aspects of our approach should be taken into consideration. Besides accurate georeferencing of all samples, the number and spatial distribution of samples should be carefully considered. As it is true for previous methods (Ramachandran *et al.* 2005 Peter & Slatkin 2013), inferences based on the GSSA will be most useful with sampling that maximizes geographical space at the cost of multiple samples per location. Samples clustered in a particular area, and representing only a small portion of the distribution of the taxon of interest, would probably carry limited information and lead to biased estimates. Similarly, our results indicate that precaution should be taken with peripheral localities as they may carry a smaller signal due to noise introduced by boundary effects. In this regard, the advantage of allowing only a single individual per locality should help researchers to more efficiently design their sampling scheme without excessive increases in overall costs, assuming that accessing most of the range of a species is not prohibitive. Furthermore, larger sampling of geographic space could potentially better enable the implementation of a spatial interpolation to identify major colonization routes in more detail (Li & Heap 2011).

Like any method using genomewide SNP data, another relevant consideration is the potential for confounded inference if the sampled SNPs are impacted from parts of the genome under strong natural selection (Sokal *et al.* 1989 Lotterhos & Whitlock 2014), even if they are not linked (Allman & Weissman 2018). For this reason, it would be advisable to test for selective neutrality *a priori*, and to remove loci potentially under selection. An additional consideration is that care should be taken to ensure that inference is not confounded by multiple population histories that involve different sources of expansion. Although use of the GSSA does not explicitly require clustering individuals into populations *a pirori* (Frichot & François 2015; Petkova *et al.* 2016), the conduction of exploratory analyses to obtain some information about population structure may help identify cases of admixture and more than one source for expansion.

As a stand-alone summary statistic, the GSSA is itself an exploratory tool to be used with other spatial genomic methods such as EEMS (Petkova *et al.* 2016), MAPS (Al-Asadi *et al.* 2018), or SpaceMix (Bradburd *et al.* 2016): it can investigate one’s data and help formulate model-based hypotheses about population history (House & Hahn 2017). Ideally, the GSSA should be incorporated into an inferential model that allows for testing historical hypotheses and for estimating relevant demographic parameters (He *et al.* 2017a).

### Future Prospects

Further evaluations are needed to assess whether our approach can also detect multiple sources of expansion (a bimodal distribution of raggedness indices could potentially suggest two colonization sources), and if it may allow us to estimate the relative timing of expansions (for instance, by using the slope of the spatial clines in the raggedness of the GSSAs; Fig. 3). Additionally, the GSSA might be used more generally to test for spatial expansion in the first place by using null simulations to determine the significance of spatial clines in the raggedness of GSSAs (Fig. 3). Specifically, the GSSA shape quantified by Harpending’s (1994) RI could be used to test for expansion in the context of a null distribution expected under an ahistorical equilibrium between distance and genetic differences (i.e. isolation by distance; IBD). Under the latter scenario, the allelic similarity is expected to be inversely correlated with distance (Wright 1943), and hence the spatial cline of raggedness indexes should not be significant. On the contrary, under a range expansion this slope is expected to be significantly different from zero. Finally, the GSSA could also be a useful way to explore a broader set of models such as a source/sink population scenarios (MartinezSolano & Gonzalez 2008), cyclical histories of admixture (Frantz *et al.* 2013 Alvarado-Serrano & Hickerson 2015), or heterogeneous population densities (Excoffier *et al.* 2008). However, the potential for these future directions remains speculative at this point and further work is needed to assess these possibilities.

Our approach may also be expanded if there is interest in more precisely identifying the geographic origin of an expansion when the source area is suspected to be missing from the sample. That would require interpolating the expected raggedness index at un-sampled localities to identify the region with the smallest indexes. Implementing an interpolation would also allow us to potentially identify spatially isolated areas of contiguous high raggedness indices, which could serve to uncover instances of expansion from more than one source using an Euclidean allocation algorithm. However, as interpolation is strongly impacted by the number and distribution of observed samples (Stein 2012), its implementation may be not possible when a limited number of locations are available since interpolation accuracy drastically decreases for small sample sizes. Alternatively, a time-difference of arrival approach (TDOA; (Gustafsson *et al.* 1994)) could be implemented, as done by Peter and Slatkin’s (2013) ψ. Commonly used for localization, the TDOA is a triangulation approach that takes advantage of the change in the strength of a signal as distance from the source increases (Gustafsson *et al.* 1994 Drake & Dogancay 2004). Specifically in our case, the pairwise difference in raggedness index between location pairs may be used as the signal change for triangulation to identify the geographic coordinates of the hypothesized source (see eq. 5 in Peter and Slatkin 2013).

For those interested in community ecology and species interactions, the GSSA may be used in aggregated geo-referenced population genomic datasets of co-distributed taxa to test for concerted spatio-temporal dynamics between two interacting species (Perkins & Swayne 2001 Wicker *et al.* 2012), the sources of multiple invading species (Sax *et al.* 2007 Johnson *et al.* 2009), and the geographic origins of multiple historic domestications (Kanginakudru *et al.* 2008 He *et al.* 2011). Likewise, it may be used to help identify shared regions of secondary contact and hybridization between long-isolated co-distributed pairs of taxa (Remington 1968 Moritz *et al.* 2009), or to understand the assembly of whole communities across geographic barriers or trajectories of expansion after global shifts in climate (Avise *et al.* 1987 Ibrahim *et al.* 1996 Avise 2000 Hewitt 2000 Burbrink *et al.* 2016). Additionally, if one were interested in incorporating a resistance surface to correct for effective distances that emerge from landscape features (Spear *et al.* 2010), the GSSA may easily accommodate alternative distance matrices to allow the integration of realistic landscape scenarios.

## Conclusion

The GSSA statistic we present here builds on emerging efforts to make phylogeography and population genetics more spatially explicit (Alvarado-Serrano & Hickerson 2015 House & Hahn 2017 Ashander *et al.* 2018 Bradburd *et al.* 2018). By doing so, it improves our ability to estimate the geographical source region (or closest areas) as well as the general direction of range expansions. The use of single samples per location, and un-polarized SNPs, makes this metric well-suited for a wide range of non-model organisms. The GSSA offers not only the capability to serve as an additional spatially explicit summary of genetic patterns for modelbased inference when likelihoodbased methods are intractable (Beaumont 2010 Pudlo *et al.* 2015), but, most importantly, presents a tool for guiding the development of spatially-explicit demographic hypotheses by better accounting for the spatial component of species histories. Further developments of this statistic, including the refinement of an interpolation procedure for geographically sparse samples, should lead to easier incorporation of this tool into simulationbased inferential approaches (Currat *et al.* 2004 Leblois *et al.* 2009 He *et al.* 2017a). We expect that the incorporation of expansion surfaces will enable more complex scenarios that include environmental heterogeneity and explicit geographic barriers to be modeled into these simulation approaches (Ray & Excoffier 2010 Joseph *et al.* 2016). Likewise, we expect clines in GSSA estimates to be able to help identify relevant spatial features, such as shared contact zones between populations that have been colonized from different expansion clusters (Swenson 2010), or consistent geographic barriers that have maintained isolation of populations during expansions and hence have promoted differentiation (Carstens *et al.* 2005 Potter *et al.* 2017). The GSSA offers a useful and flexible tool that complements existing methods for improved understanding of the processes governing population differentiation and spatial patterns of genomic diversity.

### Software

A program to calculate the GSSA and Harpending’s (1944) raggedness index from georeferenced genotypes and associated user-defined geographic distances among sampled locations is available in https://bitbucket.org/diegofalvarados/GSSA-v0.0. This software also contains the pipeline needed to recreate the simulations we used to test the accuracy of the GSSA and Peter and Slatkin’s ψ. A full step-by-step implementation of our method, including comments and visualization of the intermediate outputs, is available at the following jupyter notebook: https://mybinder.org/v2/gh/ftempo/GSSA/master.

## Acknowledgements

We would like to thank Benjamin Peter for assistance in the implementation of the directionality method and Zach Gompert for generously providing the genomewide georeferenced SNP data, as well as Gideon Bradburd, Ana Carnaval, Marcelo Gehara, Isaac Overcast, and John Robinson for insightful comments on previous versions of this manuscript. All authors reviewed the manuscript. Funding was provided by grants from FAPESP (BIOTA, 2013/50297-0 to MJH and AC Carnaval), NASA through the Dimensions of Biodiversity Program (DOB 1343578 to MJH), and the National Science Foundation (DEB-1253710 to MJH). This work would not have been possible without help from the City University of New York High Performance Computing Center, with support from the National Science Foundation (CNS-0855217 and CNS-0958379).

## Data Accessibility

The whole pipeline and associated files needed to recreate the simulations used is available at the following bitbucket repository: https://bitbucket.org/diegofalvarado-s/GSSA-v0.0

## Author Contributions

DFA and MJH conceived the GSSA statistic and DFA led the development of it. DFA coded the simulations, analyses and produced the figures. DFA and MJH wrote the manuscript.

---

**Supp. Table 1.**
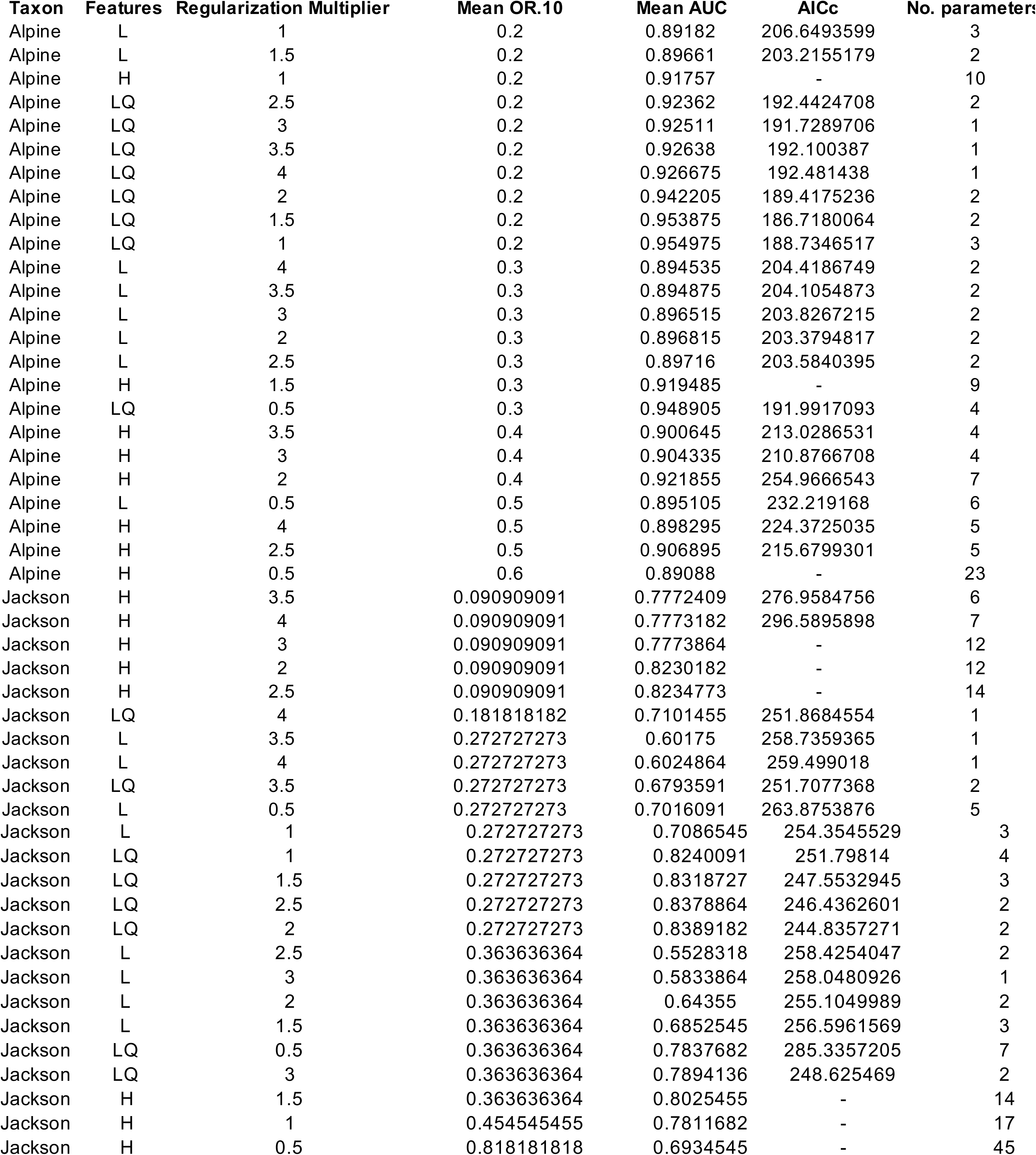
Tuning results for ecological niche models of *Lycaeides* taxa based on jacknife crossvalidation. Abbreviations: AUC: areaunder the ROC curve; OR.10: 10^th^ percentile omission rate AICc: corrected Akaike Information Criterion; L: linear features, Q: quadratic features, LQ: linearquadratic features H: hinge features. Models for each species are ordered according to their ranking, with the best model on top.

**Supp. Fig. 1.**
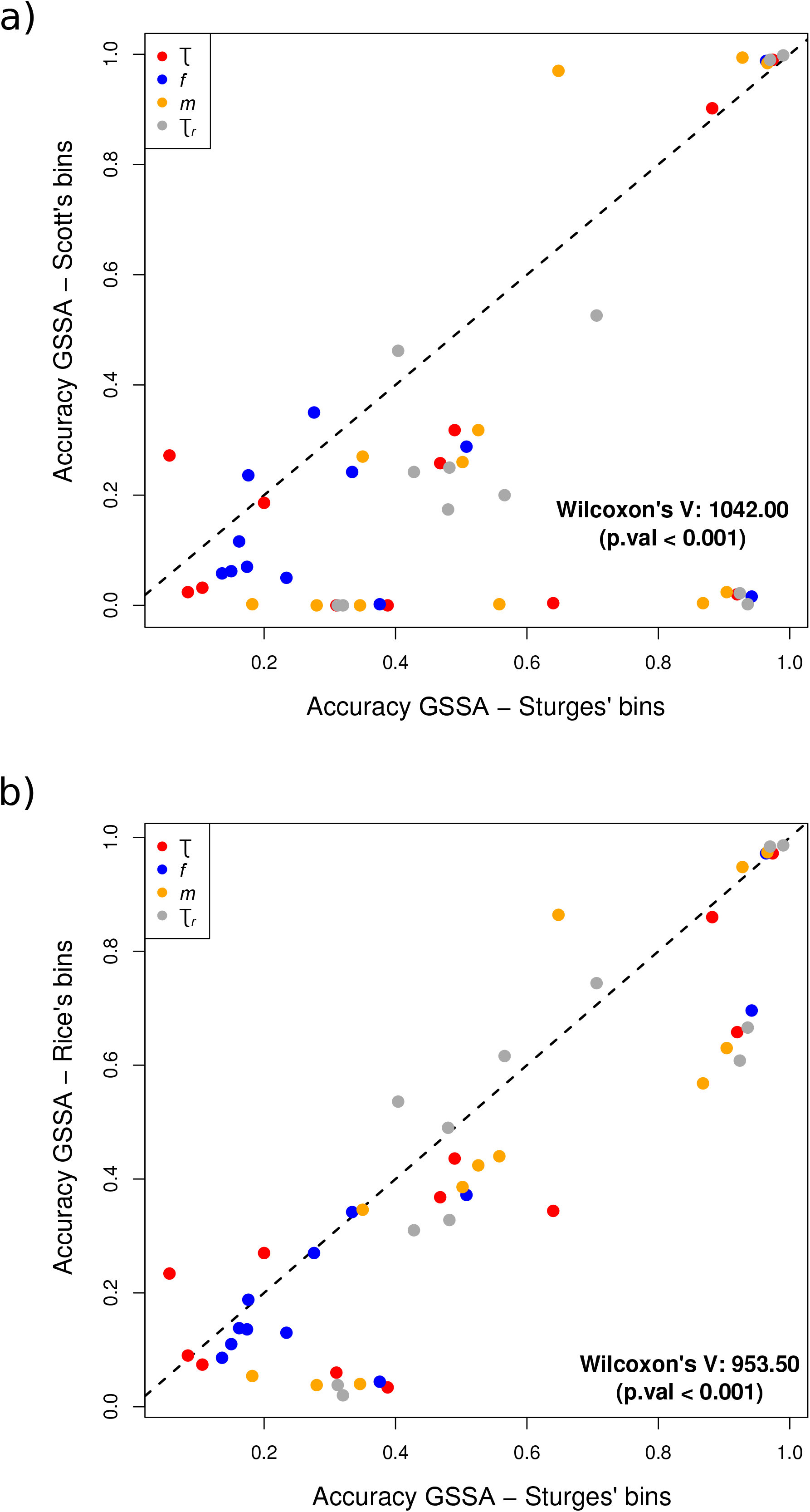
Estimation accuracy of the raggedness index calculated from the GSSA vector under alternative histogram binning schemes. Accuracy of the GSSA-based inference when constructed using Sturge’s binning equation is compared against its accuracy when constructed using a coarser (a), or a finergrain binning scheme (b). Each dot corresponds to the mean accuracy with which the source of expansion is inferred for each parameter combination and binning scheme used (points are colored according to the parameters used in the simulations see Table 1). The dashed line corresponds to a 1:1 line.

**Supp. Fig. 2.**
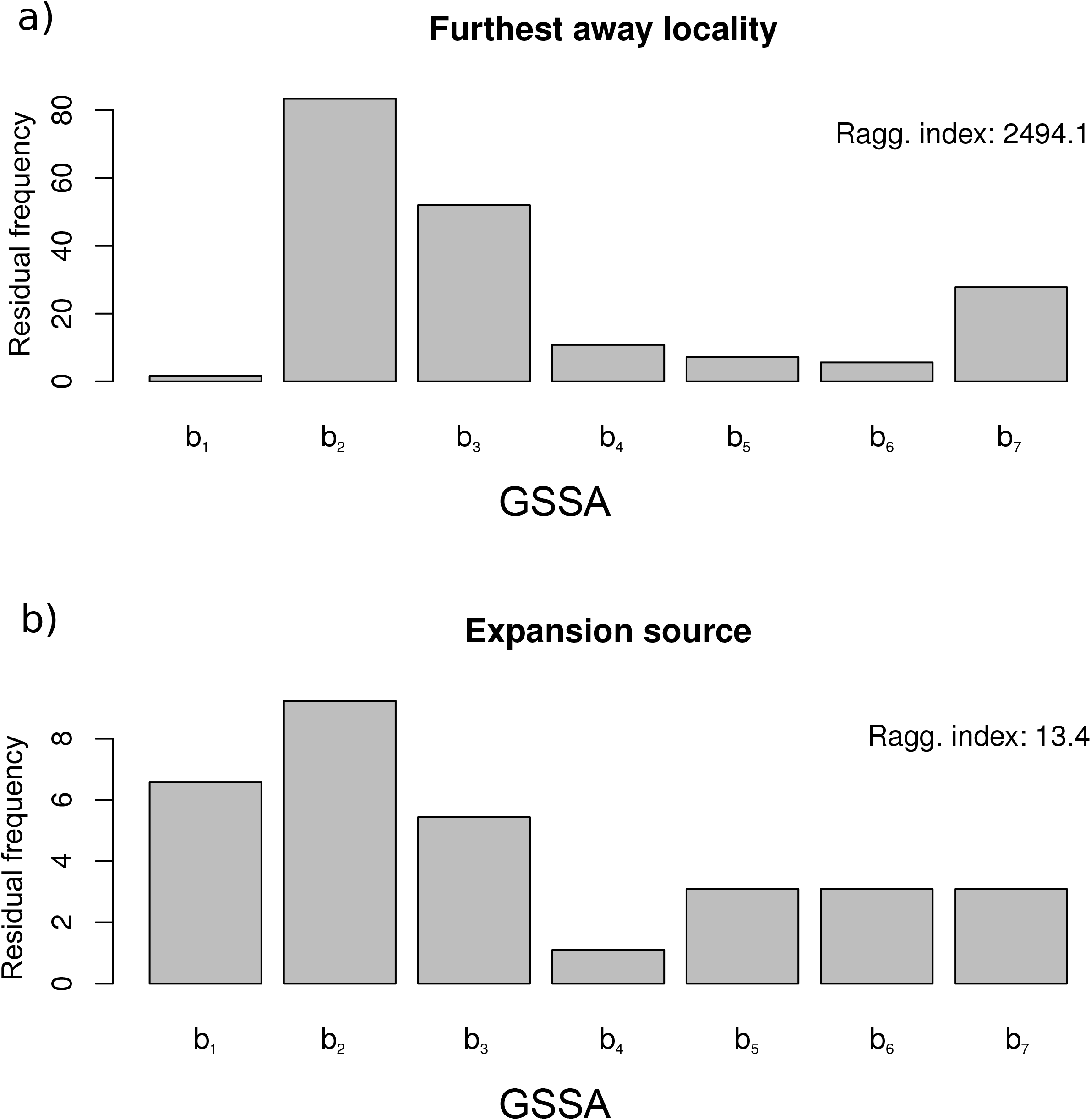
Example of GSSA vectors for (a) last-colonized and (b) the expansion source locality in one of the simulations. The frequency of each of the GSSA bin (b _k_) is depicted along_*r*_ = 0.01 *m* = the corresponding raggedness index (Simulation parameters used: *τ* = 0; *f* = 0.002; *τ* 0 *N_K_* = 1000; *τ_c_* = 0.01; simulated source = “origin B”).

**Supp. Fig. 3.**
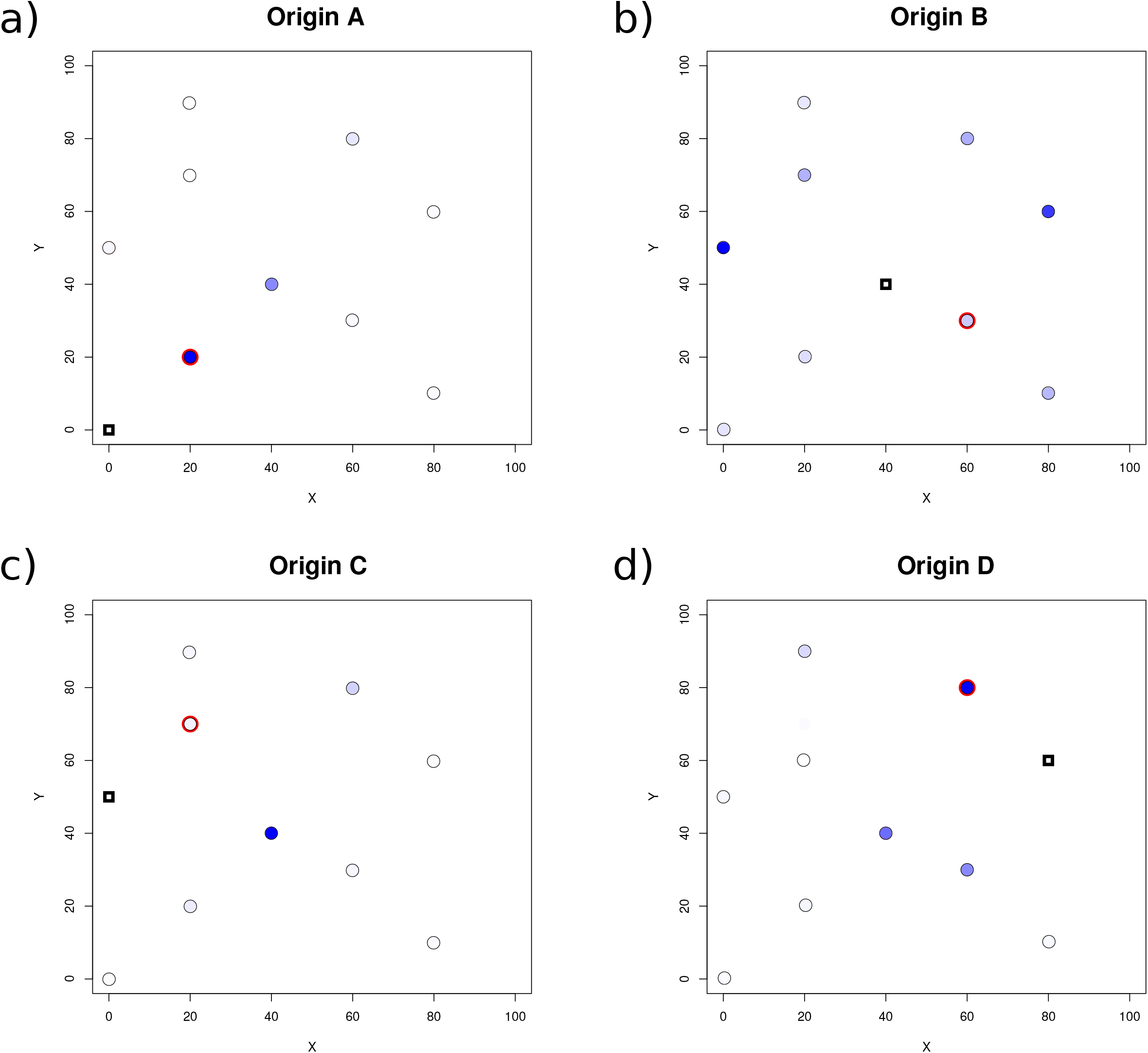
Geographic position of the sampled locality closest to the source of expansion according to the GSSA-RI approach. A red circle denotes the locality, among those sampled (indicated with black incircles), that was first colonized (note the simulated source itself, denoted by a black square, is not sampled). Results presented are aggregated across simulations, with the relative frequency that each position was identified as the first to be colonized indicated by the color hue (darker colors corresponding to greater frequencies). Note that in the vast majority of instances, the GSSA-RI approach correctly infers the closest neighbor to the source or another closeby neighbor location.

